# TIMP-1 attenuates the development of inflammatory pain through MMP-dependent and receptor-mediated cell signaling mechanisms

**DOI:** 10.1101/540724

**Authors:** B.E. Knight, N. Kozlowski, J. Havelin, T. King, S.J. Crocker, E.E. Young, K.M. Baumbauer

## Abstract

Unresolved inflammation is a significant predictor for developing chronic pain, and targeting the mechanisms underlying inflammation offers opportunities for therapeutic intervention. During inflammation, matrix metalloproteinase (MMP) activity contributes to tissue remodeling and inflammatory signaling, and is regulated by tissue inhibitors of metalloproteinases (TIMPs). TIMP−1 and −2 have known roles in pain, but only in the context of MMP inhibition. However, TIMP-1 also has receptor-mediated cell signaling functions that are not well understood. Here, we examined how TIMP-1-dependent cell signaling impacts inflammatory hypersensitivity and ongoing pain. We found that hindpaw injection of complete Freund’s adjuvant (CFA) increased cutaneous TIMP-1 expression that peaked prior to development of mechanical hypersensitivity, suggesting that TIMP-1 inhibits the development of inflammatory hypersensitivity. To examine this possibility, we injected TIMP-1 knockout (T1KO) mice with CFA and found that T1KO mice exhibited rapid onset thermal and mechanical hypersensitivity at the site of inflammation that was absent or attenuated in WT controls. We also found that T1KO mice exhibited hypersensitivity in adjacent tissues innervated by different sets of afferents, as well as skin contralateral to the site of inflammation. Replacement of recombinant murine (rm)TIMP-1 alleviated hypersensitivity when administered at the site and time of inflammation. Administration of either the MMP inhibiting N-terminal or the cell signaling C-terminal domains recapitulated the antinociceptive effect of full-length rmTIMP-1, suggesting that rmTIMP-1inhibits hypersensitivity through MMP inhibition and receptor-mediated cell signaling. We also found that hypersensitivity was not due to genotype-specific differences in MMP-9 activity or expression, nor to differences in cytokine expression. Administration of rmTIMP-1 prevented mechanical hypersensitivity and ongoing pain in WT mice, collectively suggesting a novel role for TIMP-1 in the attenuation of inflammatory pain.

**AUTHOR CONTRIBUTIONS:** BEK – experimental design, data collection and analysis, mouse breeding and care, manuscript preparation

NK – data collection

JH – collection of conditioned place preference data

TK – experimental design and analysis of data from conditioned place preference experiments, manuscript preparation

SJC-generation of T1KO mouse line, experimental design, manuscript preparation

EEY – experimental design, data analysis, manuscript preparation

KMB – experimental design, data analysis, manuscript preparation, experimental oversight

## INTRODUCTION

Tissue inhibitors of matrix metalloproteinases (TIMPs) and matrix metalloproteinases (MMPs) are released during tissue damage to facilitate tissue remodeling through degradation and reorganization of the extracellular matrix (ECM) (Gardner and Ghorpade, 2003;Nagase et al., 2006;Ries, 2014). During this process MMPs also engage an inflammatory response through proteolytic maturation of cytokines, and both of these activities are regulated through a 1:1 stoichiometric interaction with one of four tissue inhibitors of metalloproteinases (TIMP-1, −2, −3, −4) (Huang et al., 2011). The interaction between MMPs and TIMPs is tightly controlled, but research has shown that during tissue damage, imbalance between MMPs and TIMPs can lead to pathological conditions such as arthritis, multiple sclerosis, Parkinson’s Disease, cancer, and even chronic pain (Nakagawa et al., 1994;Nagase et al., 1999;Kouwenhoven et al., 2001;Yang et al., 2002;Brew and Nagase, 2010). Studies examining the role of MMPs in pain specifically have shown that increased MMP-2 and −9 activity contribute to increased pain-related behavior in response to injury that can be reversed by MMP antagonism (Kawasaki et al., 2008;Ji et al., 2009;Li et al., 2016;Remacle et al., 2018). These findings contributed, in part, to the development of several small molecule drugs that directly target and inhibit MMP activity. However, more than 50 clinical trials examining the efficacy of these drugs were discontinued due to the emergence of adverse events, including musculoskeletal pain (Cathcart and Cao, 2015;Martinho et al., 2016). While the results of these trials indicated that specific targeting of MMP activity alone is not an effective strategy for pain treatment, they also suggest that additional mechanisms related to MMP activity may contribute to pain and its inhibition, and that endogenous inhibitors of MMPs, such as TIMP-1, may attenuate pain-related behavior.

TIMP-1 is best characterized as an inhibitor of MMP activity. Indeed, TIMP-1 regulates 14 of the 24 known MMPs (Baker et al., 2002;Gardner and Ghorpade, 2003;Nagase et al., 2006;Kawasaki et al., 2008), and has been shown to prevent the development of mechanical and thermal hypersensitivity following nerve damage (Kawasaki et al., 2008;Martinho et al., 2016). However, this identified role was characterized purely in the context of MMP inhibition. TIMP-1 inhibits MMP activity through the binding of its N-terminus with the targeted MMP, resulting in chelation of Zn^2+^ from the enzyme active site (Gomis-Rüth et al., 1997). Interestingly, there is now mounting evidence that the C-terminal domain can bind to membrane bound receptors, including CD63 (Nagase et al., 2006). The binding of TIMP-1 to CD63 engages intracellular signaling events that allow TIMP-1 to function as a trophic factor and initiate cellular migration and differentiation (Gardner and Ghorpade, 2003;Jourquin et al., 2005;Thorne et al., 2009;Moore and Crocker, 2012;Claycomb et al., 2013). Because TIMP-1 and MMPs can be up-regulated simultaneously during tissue damage and repair, such as in peripheral nerve injury (Parkitna et al., 2006;Huang et al., 2011;Kim et al., 2012;Yokose et al., 2012), disentangling how TIMP-1 regulates tissue remodeling/repair from the induction of pain, *per se*, is challenging.

Inflammation is a core component of the nerve injury process (Tracey and Walker, 1995), and, in general, is a significant predictor of pain chronicity (Tal, 1999;Kehlet et al., 2006;Chapman and Vierck, 2017). Therefore, we used a model of cutaneous inflammation to examine the effects of TIMP-1 signaling on pain in the absence of frank tissue damage. We found that hindpaw injection of complete Freund’s adjuvant (CFA) induced TIMP-1 expression in keratinocytes prior to the emergence of hypersensitivity in wildtype (WT) mice, suggesting that the release of TIMP-1 at the site of inflammation prevented the development of hypersensitivity. Supporting this conclusion, we found that TIMP-1 knockout (T1KO) mice exhibited robust hypersensitivity to stimulation of tissues local and distal to the site of inflammation. This phenotype was prevented by the administration of recombinant murine (rm)TIMP-1, as well as the individual N- and C-terminal constructs at the time of CFA-injection. These results suggest that cell-signaling mechanisms may also contribute to the antinociceptive effects of TIMP-1. In addition, inflammation in WT and T1KO mice did not result in the genotype-specific activation or expression of cutaneous MMP-9 or cytokines. Finally, we found that the administration of rmTIMP-1 prevented ongoing inflammatory pain and evoked mechanical hypersensitivity in WT mice. Collectively, our data suggests that TIMP-1 regulates the algogenic properties of inflammation and that TIMP-1 may be a target for improving pain management.

## MATERIALS AND METHODS

### Animals

Experiments were conducted using 8-12-week-old (20-30 g) male WT (C57BL/6; Jackson Laboratories, Bar Harbor, ME) and T1KO mice that were group housed, and maintained in a temperature-controlled environment on a 12 hr light-dark cycle with free access to food and water. TIMP-1 knockout (T1KO) mice (Lee et al., 2005) were backcrossed onto a C57BL/6 background for greater than 13 successive generations and bred in-house as a homozygous line(Crocker et al., 2006). All studies were approved by the UConn Health Institutional Animal Care and Use Committee and treated in accordance with published NIH standards.

### Complete Freund’s Adjuvant (CFA)

To produce an acute, local inflammatory response, we subcutaneously (s.c.) injected the right hindpaw of mice with emulsified (50% in 10 μL) CFA (Sigma, St. Louis, MO). To assess primary hypersensitivity (i.e. at the site of inflammation) we administered CFA into the glabrous skin or ventral surface of the right hindpaw. Conversely, secondary hypersensitivity was assessed in skin that was adjacent or contralateral to the site of inflammation. All samples were compared to naïve controls because in a pilot experiment we found that vehicle injection alone caused increased sensitivity in T1KO mice. While this result is interesting, and suggests that subtle perturbations cause robust alterations in sensory thresholds, adding saline-treated mice confounds our ability to examine inflammatory hypersensitivity. Therefore, to interpret the effects of inflammation *per se*, naïve mice were used as comparison controls. The literature is also mixed on the use of vehicle controls in experiments using CFA, and our experiments are in line with previously published work (Allchorne et al., 2005;Jankowski et al., 2012;Imbe and Kimura, 2017).

### Recombinant Murine TIMP-1 Administration

WT and T1KO mice received injections (s.c.) of recombinant murine (rm)TIMP-1 (10 ng/μL, 10 μL; R&D Systems; Minneapolis, MN immediately following CFA injection (10 μL) into the right hindpaw. In subsequent experiments, T1KO mice received equimolar concentrations of the truncated C-terminus peptide (TIMP-1(C); 6.3kDa; Peptide 2.0 Inc., Chantilly, VA) that retains cell signaling function or the truncated N-terminus peptide (TIMP-1(N); 20 kDa; Abcam, Cambridge, UK) that retains MMP-inhibitory function and no cell-signaling ability, immediately following CFA injection.

### von Frey testing

All mice were place into transparent Plexiglas chambers (radius = 32 mm, height = 108 mm) on an elevated mesh screen and were allowed to acclimate for a minimum of 1 hr before testing. To assess mechanical sensitivity, the plantar surface of the right hindpaw was stimulated using von Frey filaments using the up-down method previously described by (Dixon, 1980). Nocifensive responses were counted as robust flexion responses, paw shaking, or paw licking and subtracted from individual baseline threshold to account for inter-subject variability. Data are presented as paw withdrawal thresholds (PWT; in grams).

### Thermal hyperalgesia

Thermal hyperalgesia to radiant heat was assessed using a Hargreaves apparatus (Harvard Apparatus; Holliston, MA) (Hargreaves et al., 1988). Briefly, all mice were placed in transparent Plexiglas chambers (radius = 32 mm, height= 108 mm) on top of a framed glass panel and were allowed to acclimate for a minimum of 1 hr before testing. Following the acclimation period, an infrared (IR) beam was aimed at the plantar surface of each hindpaw in an alternating fashion. The intensity of the IR beam was chosen to produce average baseline paw withdrawal latency (PWL) of 15-20 seconds. Stimuli were presented 5 times in an alternating fashion between each hindpaw with 5 minute intervals between successive stimulus exposures. A 30 seconds exposure cutoff was employed to prevent tissue damage. PWLs collected from each paw were then averaged and analyzed.

### Conditioned Place Preference (CPP)

CPP was used to assess ongoing pain in WT mice. A 3-day single trial protocol was used. On day 1, all mice freely explored a 3-chamber CPP box for 15 minutes prior to injection with CFA or saline. Preconditioning (baseline) behavior was analyzed using automated software (ANYMAZE, Stoelting) to ensure there were no baseline differences in the time spent in any of the chambers. On day 2 (conditioning day), all mice received an intrathecal (i.t.) injection of saline (5 μL volume) under isoflurane anesthesia and upon waking (within 2 min) were confined into the pre-assigned pairing chamber for 30 min. They were then returned to their home-cages for 4 hr. All mice then received an intrathecal (i.t.) injection of clonidine (2 μg/uL; 5 μL volume) through lumbar puncture and upon waking were confined to the opposite pairing chamber for 30 min. Vehicle and clonidine paired chambers were randomly assigned and counterbalanced between animals. On day 3 (test day), mice were returned to the CPP apparatus and allowed to freely explore all chambers across 15 min. The total time spent in each chamber was assessed using automated software (ANYMAZE). Conditioning day was 20-24 hr following CFA injection as previous research indicates ongoing pain is observed at this time-point (Okun et al, He et al). A total of 17 mice were used, 8 were treated with rTIMP-1 and 9 with vehicle.

### Tissue collection

Mice were anesthetized with a lethal dose of ketamine and xylazine mixture (90/10 mg/kg, respectively) and intracardially perfused with ice cold 0.9% saline prior to the dissection of ipsilateral hairy skin, L2 -L3 DRG, and L2 -L3 spinal cord segments. Tissues were collected following the completion of behavior or at designated time-points for molecular analysis.

### Enzyme-linked immunosorbent assay (ELISA)

Protein was extracted through homogenization in ice-cold RIPA buffer/protease inhibitor cocktail and spun for 20 min at 4°C at 18,000 rcf. Each sample’s total protein concentration was determined using Pierce BCA Protein Assay Kit (ThermoFisher Scientific, Waltham, MA). The following ELISAs, TIMP-1 (R&D Systems, Minneapolis, MN), MMP-9 (R&D Systems; Minneapolis, MN), IL-6 (Invitrogen Carlsbad, CA), TNF-α (ThermoFisher Scientific, Waltham, MA), IL-10 (ThermoFisher Scientific, Waltham, MA), and IL-1β (R&D Systems, Minneapolis, MN) were run according to manufacturer’s instructions. All samples were run in duplicate and absorbance ratios were read at 450nm.

### Immunohistochemistry (IHC)

Hairy skin, glabrous skin, and DRG from WT mice were excised and incubated in 0.06% brefeldin A (BFA) in serum free Hank’s balanced salt solution (HBSS) for 20 min at room temperature. Half of the samples were incubated in inflammatory soup (IS) (10uM; bradykinin triacetate, histamine dihydrochloride, serotonin hydrochloride, prostaglandin E2 dissolved in normal cerebral spinal fluid, pH 6.0) or serum-free media (Kessler et al., 1992;Hachisuka et al., 2016). Incubation in IS or serum-free media occurred for 24 hr (Kessler et al., 1992). Spinal cords from inflamed or naïve WT mice were isolated 24 hr following CFA treatment or from designated naïve controls. Prior to tissue collection, mice were intracardially perfused with 0.06% BFA in 0.9% saline for 20 min and then perfused with 4% paraformaldehyde. All samples were post-fixed overnight in 4% paraformaldehyde (PFA), cryoprotected overnight in 30% sucrose, and later embedded in Optimal Cutting Temperature (OCT). Samples were cut into 30 μm cross sections using a cryostat. Tissue sections were briefly washed with sterile phosphate buffered saline (PBS) and incubated with staining buffer (0.05% triton and 30% fetal bovine serum in PBS) solution for 40 min at room temperature. Slices were then incubated with primary unconjugated antibodies for 48hr at 4°C. The following primary antibodies were diluted in staining buffer: monoclonal anti-mouse cytokeratin 14 (K14; Abcam, Cambridge, United Kingdom; 1:300 dilution), polyclonal anti-goat TIMP-1 (R&D Systems; Minneapolis, MN; 1:300 dilution), monoclonal anti-mouse microtubule-associated protein 2 (MAP2; Millipore Sigma, Burlington, MA; 1:1000 dilution), and monoclonal anti-mouse primary conjugated-cy3 glial fibrillary acidic protein (GFAP; Abcam, Cambridge, United Kingdom; 1:500 dilution). Tissue slices were then incubated with secondary antibodies for 2-3 hr at 4°C. The following secondary antibodies were diluted in staining buffer: polyclonal rabbit anti-mouse Alexa-488 (Life Technologies, Carlsbad, CA, 1:000), polyclonal donkey anti-goat Alexa-568 (Life Technologies, Carlsbad, CA,1:000), and polyclonal goat anti-mouse Alexa-568 (Life Technologies, Carlsbad, CA; 1:1000 dilution). Slides with DRG and spinal cord slices were incubated with 300um DAPI prior to cover-slipping to visualize nuclei of satellite glial cells and astrocytes, respectively.

### MMP-9 colorimetric activity

Protein was extracted from the hindpaw of WT and T1KO mice 1 day post-CFA injections (s.c., 10μL) or from designated naïve controls for a high throughput screening of MMP-9 activity. Gelatinase activity was measured using the SensolyteGeneric MMP colorimetric assay kit (Anaspec, Fremont, CA). Samples were run in duplicate and end-point enzymatic activity was analyzed using a glutathione reference standard.

### RT-qPCR

Total RNA from WT and T1KO was extracted from naïve and inflamed samples 1 day post CFA injection using a RNeasy Mini Kit (Qiagen, Venlo, Netherlands). To quantify cutaneous TIMP-2 and TIMP-4 mRNA expression, equal amounts of cDNA were synthesized using the iScript cDNA Synthesis Kit (Bio-Rad Laboratories, Hercules, CA) and mixed with SsoAdvanced Universal SYBR Green Supermix (Bio-Rad Laboratories, Hercules, CA) and 2uM of both forward and reverse primers (see Table 1). GAPDH was amplified as an internal control. The threshold crossing value was noted for each transcript and normalized to the internal control. The relative quantitation of each transcript was performed using the ΔΔCt method and presented as fold change relative to naïve WT expression.

**Table 1:**
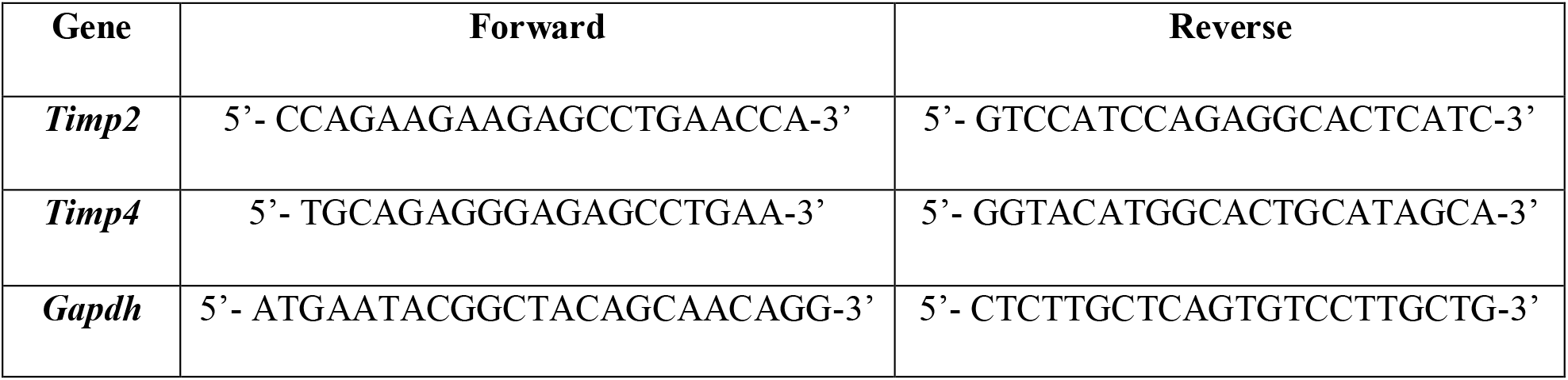
Primer sequences for qPCR.

### Statistical analysis

All data were analyzed using one-way or mixed designs Analysis of Variance (ANOVA). *Post hoc* analyses were performed using Tukey’s HSD, and statistical significance was determined using a *p*<.05. Statistical analysis was performed using SPSS (Version 25). Since ANOVAs rely on linear relationships among data, and not all effects can be resolved using linear based statistical tests, we used trend analyses (e.g., contrasts) to test for significant nonlinear relationships in some of our behavioral analyses. An added benefit of this approach is that trend analyses are more robust than ANOVAs (Tabachnick and Fidell, 2007). Clonidine induced CPP was assessed using a 2-way repeated measures ANOVA with post-hoc analysis of pre vs post conditioning time spent in the clonidine paired chamber for each treatment group using Sidak’s multiple comparisons test. Between groups analysis was performed on difference scores calculated as post-conditioning (-) pre-conditioning time spent in the clonidine paired chamber.

## RESULTS

### Cutaneous TIMP-1 expression is upregulated prior to the onset of inflammatory hypersensitivity

To determine whether cutaneous inflammation alters the expression of TIMP-1 in tissues along the peripheral sensory circuit, we injected emulsified CFA (10 μL, s.c.) into the hairy skin of the ipsilateral hindpaw and collected spinal cord (SC; L2-L3), dorsal root ganglia (DRG; L2-L3), and hairy skin over the course of 7 days. We found that inflammation did not alter the overall expression of TIMP-1 protein in SC or DRG, all *Fs* > 1.13, *p*>.05 (Figure 1 A, B). However, we observed a significant increase in cutaneous TIMP-1 protein 1, 3, 5, and 7 days following CFA administration, *F*(4,19) = 37.54, *p*<.01 (Figure 1C). To confirm the above results, and to localize the cellular source of TIMP-1 expression, immunohistochemistry (IHC) was performed on DRG and skin samples incubated *in vitro* with or without inflammatory soup (IS)(Kessler et al., 1992), as well as spinal cords following *in vivo* inflammation. Although overall TIMP-1 expression levels were unaltered in the spinal cord and DRG following inflammation, we found that TIMP-1 co-localized with glial fibrillary acidic protein (GFAP) expressing cells following inflammatory stimulation, demonstrating that astrocytes (Figures 2A) and satellite glial cells (Figure 2B) appear to express TIMP-1 during inflammation (Huang et al., 2011;Welser-Alves et al., 2011). We also found that TIMP-1 expression was upregulated in K14-positive keratinocytes in both hairy and glabrous skin following inflammatory stimulation (Figure 2C; glabrous skin data not shown).

**Figure 1.**
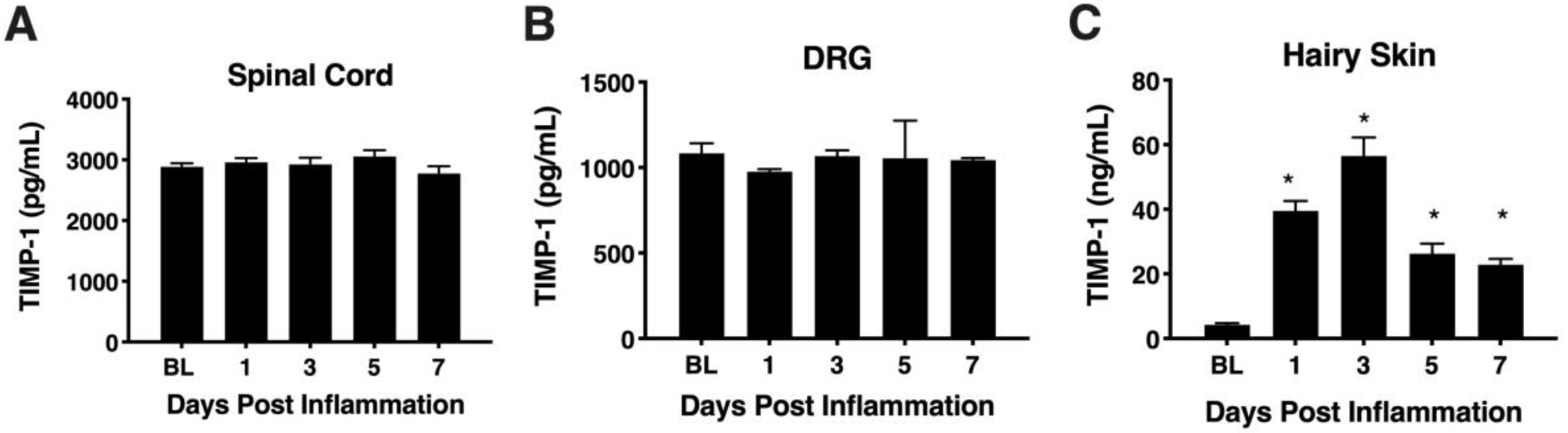
Assessing TIMP-1 expression along peripheral nociceptive circuit following cutaneous inflammation. (A) Cutaneous inflammation does not alter overall TIMP-1 protein expression in lumbar spinal cord or (B) DRG, but does increase protein expression in (C) hairy skin. n=4/condition, * indicate significant differences compared to naïve controls, *p*<.05, and error bars depict SEM.

**Figure 2.**
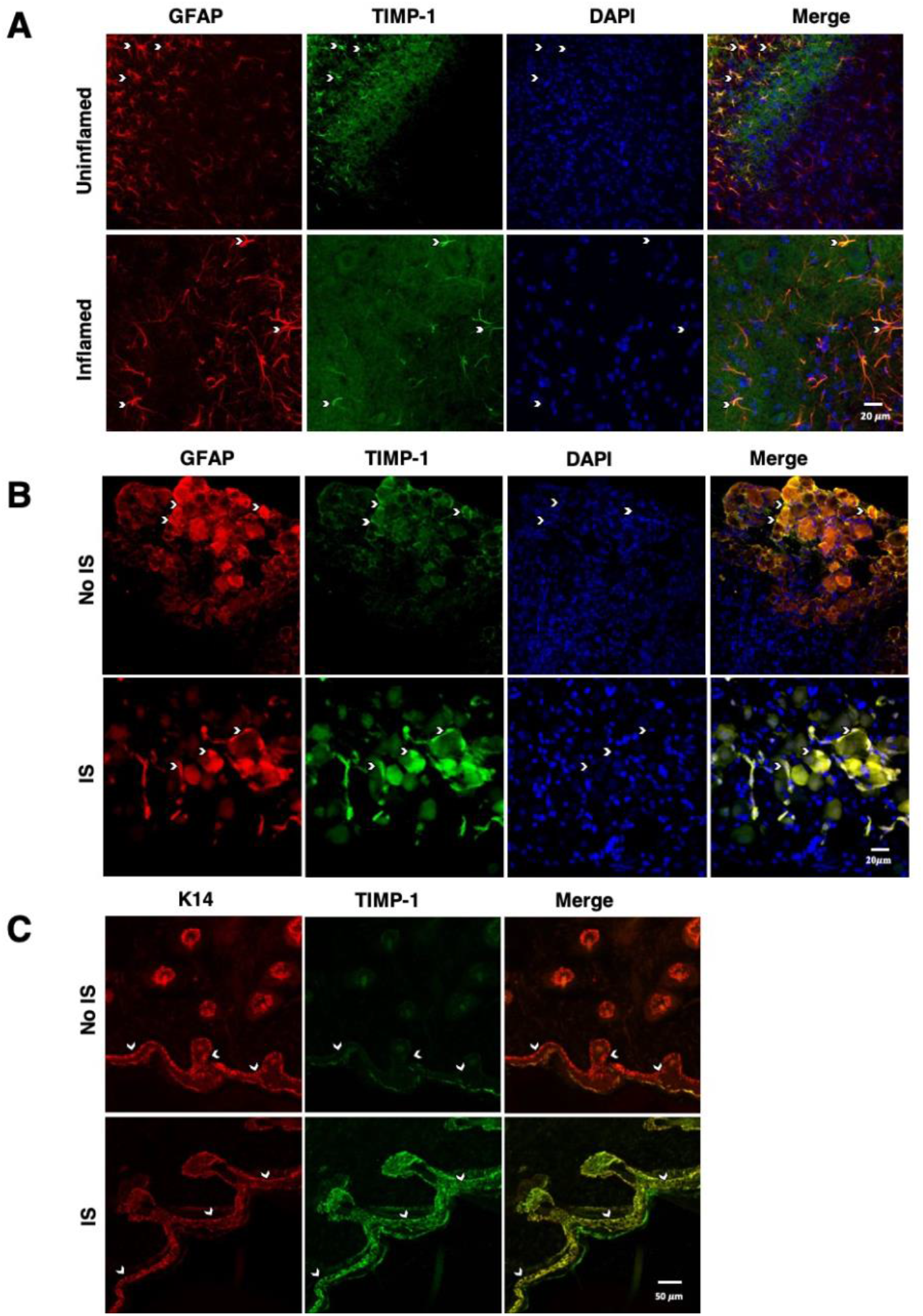
Cellular colocalization of TIMP-1 expression. (A) Immunostaining (20X) of naïve and inflamed lumbar spinal cord 24 hr following inflammation. TIMP-1 (green) expression is localized to GFAP-positive astrocytes (red). n=3/condition, scale bar 20 μm. (B) Immunostaining (20X) of naïve and inflamed lumbar DRG 24 hr following inflammation. TIMP-1 (green) expression is colocalized with by GFAP-positive satellite glial cells (red). n=3/condition, scale bar 20 μm. (C) Immunostaining (40X) of hindpaw hairy skin shows K14-positive keratinocytes (red) upregulate TIMP-1 (green) 24hr following inflammation compared to naïve control. n=3/condition, scale bar 50 μm.

To associate the expression of cutaneous TIMP-1 with the development of mechanical hypersensitivity, we assessed paw withdrawal thresholds (PWT) on the plantar surface of the hindpaw for 7 days following CFA injection into the dorsal, hairy skin. We found that TIMP-1 protein levels peaked 3 days following CFA administration (see Figure 1C), at a time when mice developed mechanical hypersensitivity, *F*(2,12) = 43.94, *p*< 0.05, (Figure 3). Together, these data indicate that cutaneous inflammation induces the expression of TIMP-1 in keratinocytes at the time of inflammation and prior to the onset of mechanical allodynia.

**Figure 3.**
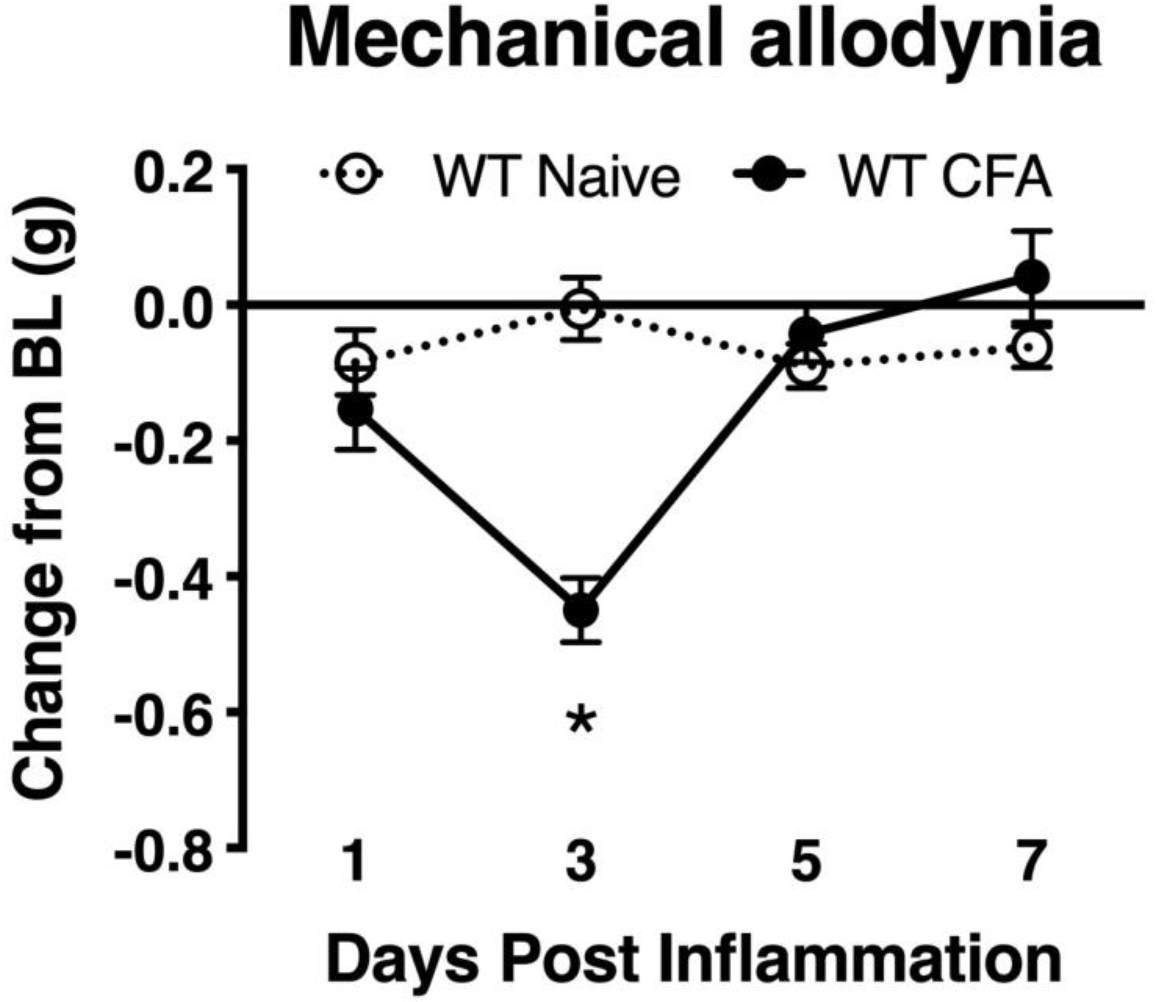
Development of inflammatory hypersensitivity in WT mice. Assessment of mechanical hypersensitivity over 7 days following s.c. administration of CFA. Inflamed mice show a significant reduction in mechanical thresholds relative to naïve mice 3 days following inflammation (n=6/condition). * indicate significant differences compared to naïve controls, *p*<.05, and error bars depict SEM.

### Mice lacking TIMP-1 exhibit hyperalgesia in inflamed and uninflamed cutaneous tissues

To determine whether endogenous TIMP-1 expression is important for the normal progression of hypersensitivity, we used a global TIMP-1 knockout (T1KO) mouse strain. We first assessed behavioral responsiveness to radiant heat on the plantar surface of the hindpaw following s.c. administration of a diluted CFA solution. We chose to use a diluted CFA solution because our preliminary experiments suggested that exposure to slight challenges significantly altered sensitivity in T1KO. To ensure that any potential differences in responding to inflammatory stimulation were not due to preexisting differences in sensory thresholds between mouse strains, we measured baseline responding to radiant heat and found no significant differences in paw withdrawal latencies (PWL), *F*(1, 31) = .47, p>.05 (Figure 4A). Interestingly, while we did not observe any significant differences in PWL between naïve and WT mice that received diluted (e.g., subthreshold) CFA, we did find that inflamed T1KO mice exhibited thermal hyperalgesia that persisted for 29 days in response to diluted CFA injection, all *Fs* < 2.30, *p* < .05 (Figure 4B).

**Figure 4.**
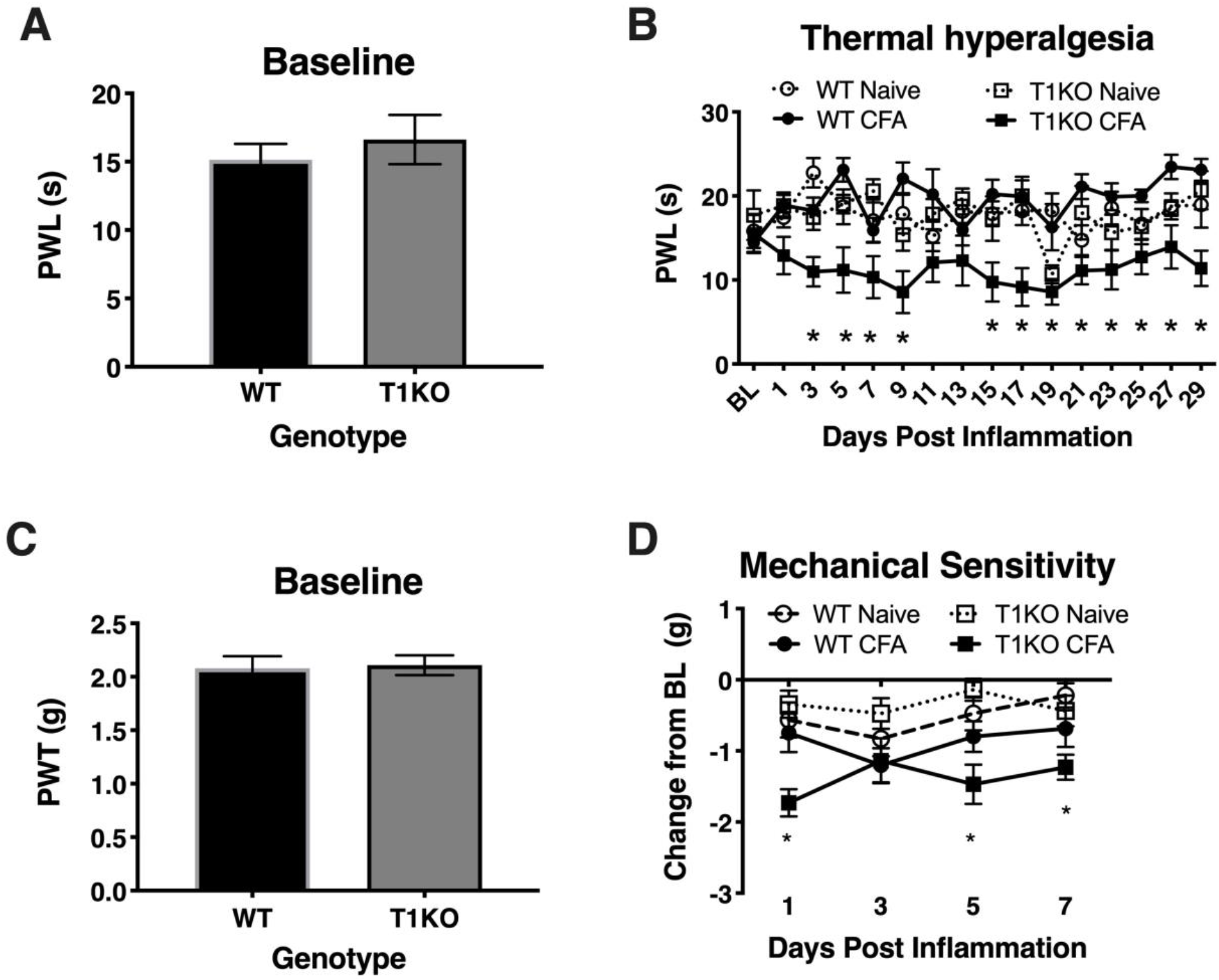
Mice lacking TIMP-1 develop thermal and mechanical hypersensitivity following cutaneous inflammation. (A) No differences in baseline thermal PWL are exhibited between T1KO and WT mice (n=16/condition). (B) Inflamed T1KO mice exhibit significantly reduced PWLs compared to inflamed WT mice and naïve WT and T1KO mice (n=8/condition). (C) Baseline assessment of mechanical PWTs revealed no genotypic differences between T1KO (n=20) and WT mice (n=18). (D) T1KO mice develop significantly reduced PWTs 1, 5, and 7 days following CFA administration compared to WT controls (n=8-10/condition).

Next, we assessed mechanical response thresholds (von Frey) following diluted CFA administration. Analysis of baseline responses to mechanical stimulation did not reveal any significant differences between genotypes *F*(1, 36) = .34, p < .05 (Figure 4C). We did find that CFA administration reduced mechanical response thresholds in both genotypes *F*(1, 34) = 17.61, *p* < .01. However, T1KO mice exhibited greater mechanical hypersensitivity 1 day following CFA treatment, compared to WT controls, all *Fs* < 4.59, p <.05 (Figure 4D). Therefore, we concluded that the up-regulation of TIMP-1 following inflammation delays the onset of hypersensitivity, and that in the absence of TIMP-1, the normal development of hypersensitivity is altered.

To determine whether the rapid emergence of inflammatory hypersensitivity in T1KO mice was due to compensatory expression of *Timp2* or *Timp4* mRNA, we examined cutaneous expression of each transcript in naïve and inflamed T1KO mice. We found no significant differences in the basal expression of either transcript in WT or T1KO mice. However, *Timp2* and *Timp4* expression decreased in T1KO mice following inflammation (Figure 5A, B), suggesting that genetic deletion of TIMP-1 did not result in a compensatory response from other TIMPs, all *Fs* < 11.65, *p* < 0.01.

**Figure 5.**
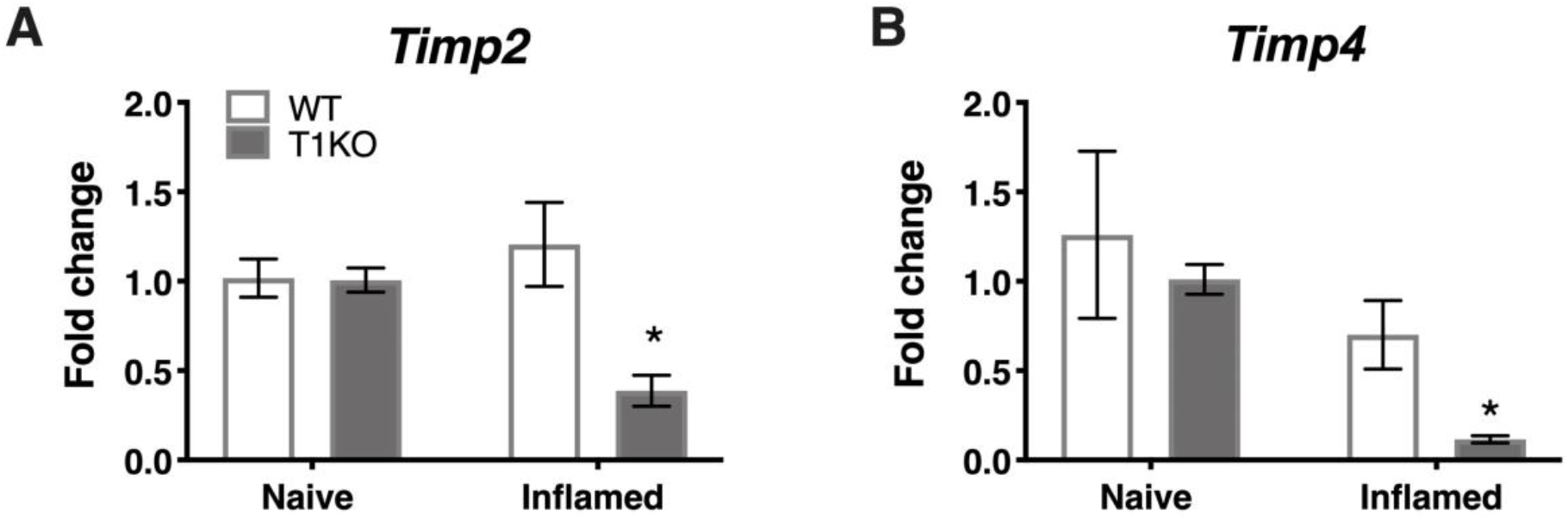
Assessment of cutaneous *Timp2* and *Timp4* mRNA expression following inflammation. (A) *Timp2* mRNA expression is decreased in T1KO mouse skin 1 day following inflammation relative to WT controls. (B) *Timp4* mRNA expression is decreased 1 day following CFA compared to WT inflamed mice. n=4/condition, * indicate significant differences compared to naïve controls, *p*<.05, and error bars depict SEM.

Our current data demonstrate that cutaneous TIMP-1 is an early emergent protein following inflammation, and that the absence of TIMP-1 alters the normal development of hypersensitivity. Therefore, TIMP-1 signaling may have important implications for regulating the development of inflammatory hypersensitivity in tissue adjacent to the site of inflammation that is innervated by afferent terminals that are different from those that innervate inflamed skin. To test this possibility, we assessed the development of hypersensitivity in the glabrous skin following injection of diluted CFA into hairy skin in both T1KO and WT following baseline assessment of sensitivity. Again, we observed no genotype-specific differences in baseline reactivity, and because of this consistent finding, we will no longer present data depicting baseline behavioral reactivity. Analysis using an ANOVA revealed that T1KO mice, relative to WT mice, exhibited increased sensitivity to mechanical stimulation on the plantar surface of the hindpaw following inflammation of hairy skin that was not temporally dependent, all *Fs* < 4.41, *p* <.05, (Figure 6A, B). However, trend analyses revealed that inflamed T1KO mice exhibited increased inflammatory hypersensitivity 1 day following CFA treatment when compared to all other mice, *F*(1,57) = 11.55, *p* <.01 (Figure 6A).

**Figure 6.**
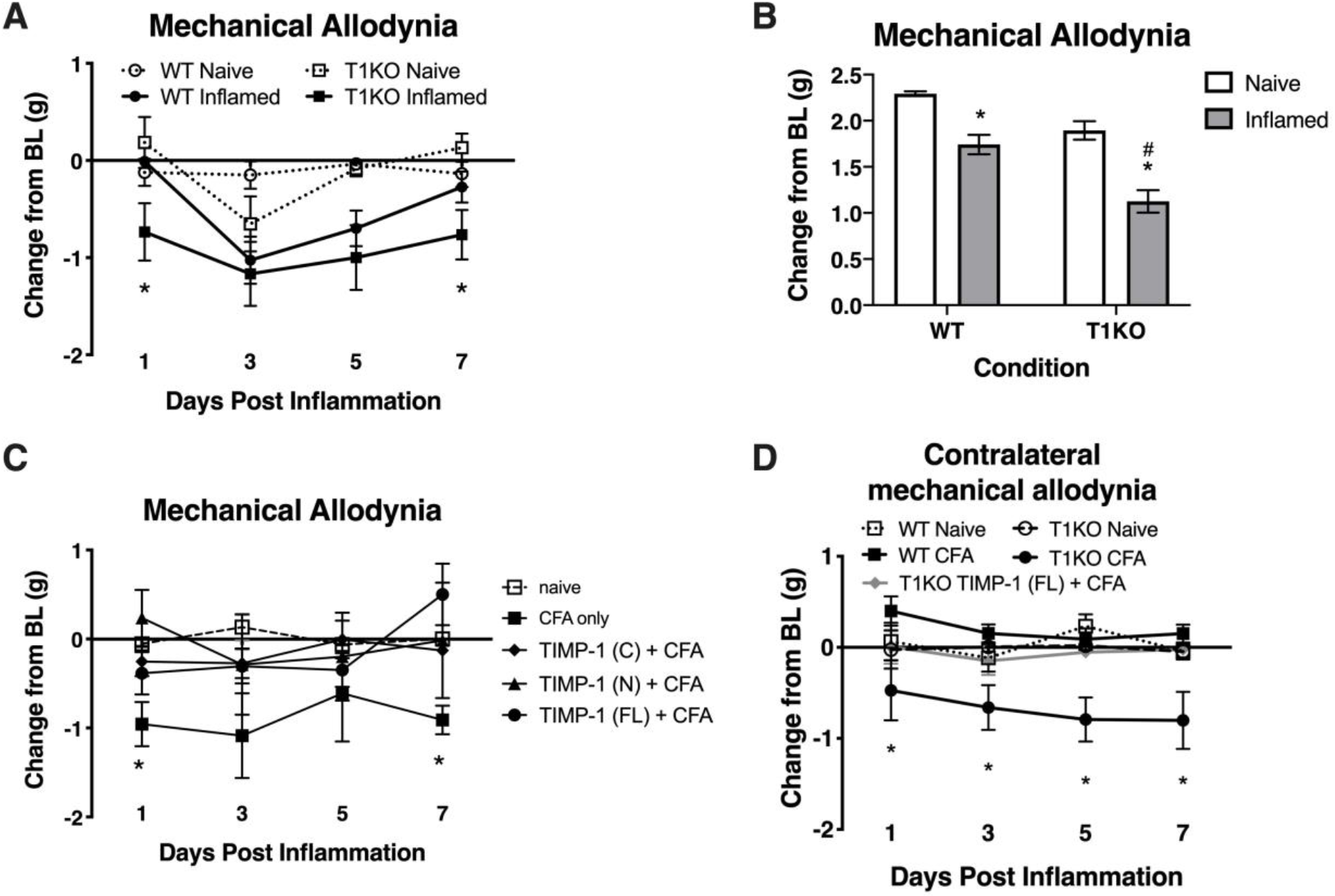
Mice lacking TIMP-1 show increased sensitivity in non-inflamed tissues. (A) Injection of CFA into the hairy skin causes mechanical hypersensitivity on the plantar surface of the paw to develop 1 day following inflammation in T1KO, but not WT, mice. (B) Graph depicting mechanical responsiveness following inflammation collapsed across time. Inflamed T1KO mice greater mechanical sensitivity overall following cutaneous inflammation. (C) Administration of TIMP-1(FL), TIMP-1(N), or TIMP-1(C) into the hairy skin at the time of inflammation prevents the development of mechanical hypersensitivity in T1KO mice. (D) Hindpaw administration of CFA produces mechanical hypersensitivity on the paw contralateral to inflammation in T1KO relative to WT mice. Treatment with rmTIMP-1 attenuated contralateral hypersensitivity in T1KO mice. PWT are presented as change from baseline. n=8/condition, * represent significant differences relative to naïve controls, *p*<.05, and error bars depict SEM.

To examine whether administration of recombinant murine (rm)TIMP-1 prevented inflammatory hypersensitivity, and the potential mechanism by which this effect occurs, separate groups of T1KO mice received a single injection of recombinant full-length rmTIMP-1 [TIMP-1(FL)], the truncated N terminus peptide [TIMP-1(N)] that retains MMP inhibitory function but no cell signaling capacity, or the truncated C terminus peptide [TIMP-1(C)]) that lacks MMP inhibitory capacity but retains its cell signaling function at the time of CFA administration. To limit the complexity of our experimental design, and to determine the optimal dose for the administration of each TIMP-1 construct, we conducted a pilot experiment using a small cohort of T1KO mice given 1, 10, or 100 ng/μL of TIMP-1 at the time of inflammation. We found that 10 ng/μL was effective at reducing inflammatory hypersensitivity (data not shown). We then administered a separate cohort of T1KO mice 10 ng/μL of TIMP-1(FL), TIMP-1(N), or TIMP-1(C) at the time of CFA administration. Mechanical hypersensitivity was assessed 24 hr later. While inflamed T1KO mice exhibited a significant reduction in mechanical thresholds, T1KO mice treated with the rmTIMP-1 peptide constructs did not. Moreover, we observed no significant differences in the response thresholds between mice given TIMP-1(FL), TIMP-1(N), or TIMP-1(C), all *Fs* > 4.54, p < .01 (Figure 6C), demonstrating that TIMP-1 attenuates inflammatory hypersensitivity through MMP-dependent and MMP-independent signaling mechanisms.

The above data show that inflammation in one somatic region could lead to mechanical hypersensitivity in tissue distal to the site of inflammation, reminiscent of “mirror image pain” (Treede et al., 1992;Treede et al., 2015). To test this possibility, we inflamed one hindpaw and measured mechanical sensitivity on the opposite hindpaw for 7 days following CFA administration. We also examined whether treatment with rmTIMP-1 at the site and time of inflammation affected sensitivity. We found that inflamed T1KO mice exhibited contralateral mechanical hypersensitivity over the course of 7 days following CFA-injection relative to WT mice (Figure 5D). Interestingly, this contralateral hypersensitivity was prevented by treatment with rmTIMP-1 in T1KO mice, *F* (4, 43) = 5.52, *p* < 0.05 (Figure 6D).

### The lack of TIMP-1 does not alter the expression of local inflammatory molecules

TIMP-1 is primarily known as a broad-spectrum MMP inhibitor, and because MMPs are known to contribute to hypersensitivity, we hypothesized that the absence of TIMP-1 may cause hypersensitivity due to elevated activity and expression of cutaneous MMP-9 (Kawasaki et al., 2008;Brew and Nagase, 2010). Examination of hairy skin collected 1 day following CFA from WT and T1KO mice demonstrated that there was an inflammation-induced increase in both MMP-9 expression and activity, all *Fs* > 7.61, *p* <.05 but that these effects were not genotype-specific, all *Fs* < 2.05, *p* >.05 (Figure 7A, B). The TIMP/MMP axis also regulates the proteolytic maturation of inflammatory molecules which can cause hypersensitivity (Pagenstecher et al., 1998). We next assessed whether the absence of TIMP-1 during inflammation caused elevated cytokine expression in the skin. Using ELISAs, we assessed the expression of cutaneous IL-1β, IL-6, TNF-*α*, and IL-10 at 1 day following CFA-injection. Analysis revealed an inflammation-induced increase in IL-1β and IL-6 expression, but this increase in expression was not different between genotypes, all *Fs* > 12.94, *p* <.05 (Figure 7C, D). In comparison, analysis of TNF-α and IL-10 did not reveal any significant differences following inflammation, all *Fs* < 4.64, *p* > .05 (Figure 6E, F). These data suggest that TIMP-1 does not affect the emergence of hypersensitivity through differences in inflammatory cytokine expression.

**Figure 7.**
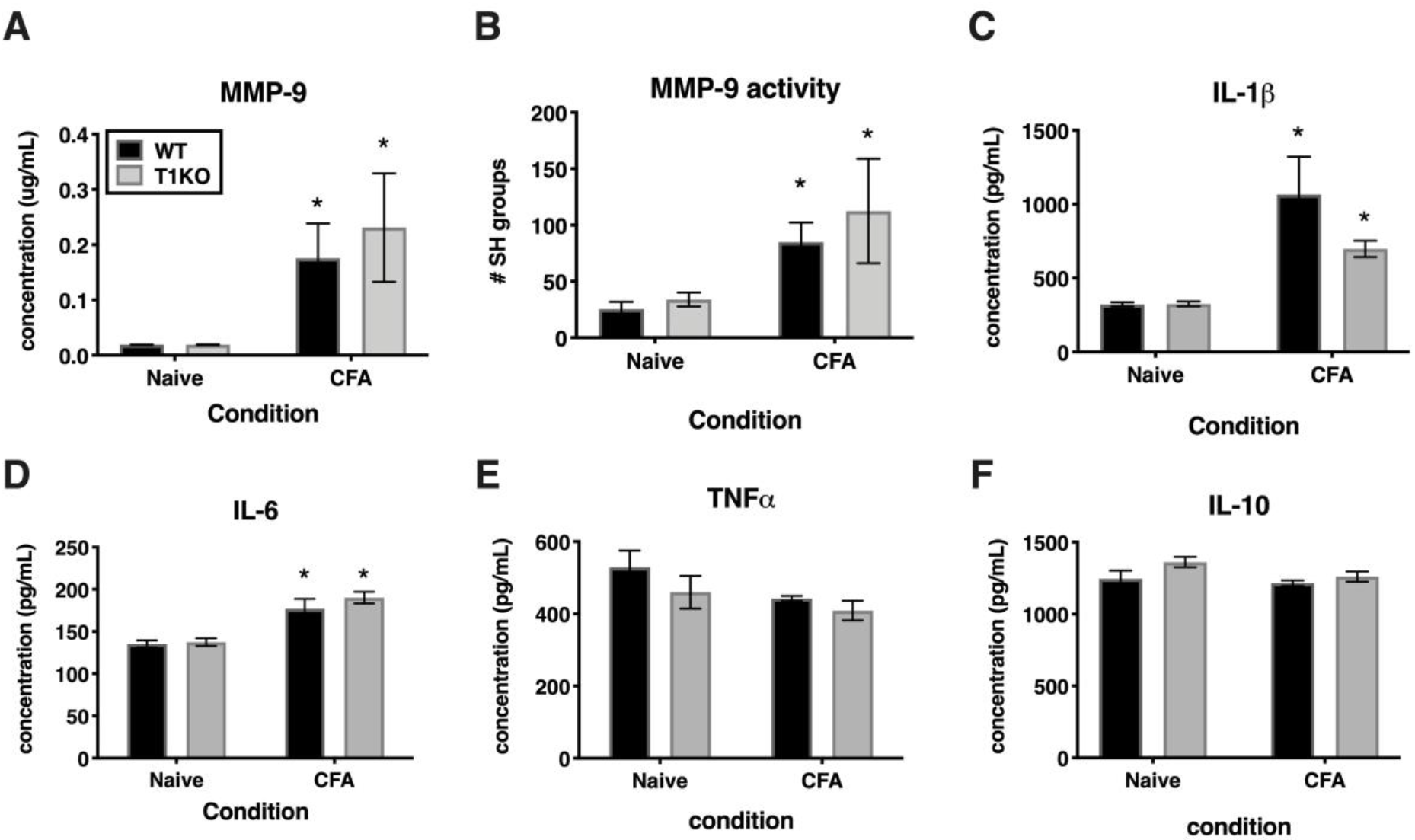
Inflammation does not alter pro-inflammatory molecules in a genotype-specific manner. A) Cutaneous inflammation significantly increases MMP-9 protein expression in WT and T1KO skin 1 day following CFA administration (n=7/condition). B) Cutaneous inflammation increases MMP-9 activity in WT and T1KO hairy skin 1 day following CFA administration. C) Cutaneous inflammation significantly increases IL-1β protein expression in WT and T1KO hairy skin 1 day following inflammation. D) Cutaneous inflammation significantly increases IL-6 protein expression in WT and T1KO hairy skin 1 day following CFA administration. E) Cutaneous inflammation does not affect expression of TNF-α following CFA administration. F) Cutaneous inflammation does not affect expression of IL-10 protein in WT and T1KO skin following CFA administration. n=4/condition, * represent significant differences relative to naïve controls, *p*<.05, and error bars depict SEM.

### Administration of recombinant TIMP-1 attenuates ongoing pain in WT mice

Previous experiments demonstrate that the administration of rmTIMP-1 attenuates evoked mechanical and thermal hypersensitivity in T1KO mice. Here, we examined whether the administration of rmTIMP-1 also attenuated ongoing pain in WT mice using CPP as previous described (King et al., 2009;Ellis and Bennett, 2013)). Analysis of pre-compared to post-conditioning time spent in the conditioning chamber indicate an effect of drug, *F*(1, 30) = 7.269, *p*<0.05), and post-hoc analysis confirmed an increase in post-conditioning time spent in the clonidine paired chamber compared to pre-conditioning time in vehicle treated mice (*p*<0.01) but not in rmTIMP-1 treated mice (*p*>0.05) (Figure 8). These observations indicate that WT mice administered CFA and TIMP-1 did not demonstrate clonidine-induced CPP (Figure 8). Because clonidine only produces CPP in the state of injury (King et al., 2009), these results further suggest that treatment with rmTIMP-1 attenuated ongoing inflammatory pain.

**Figure 8.**
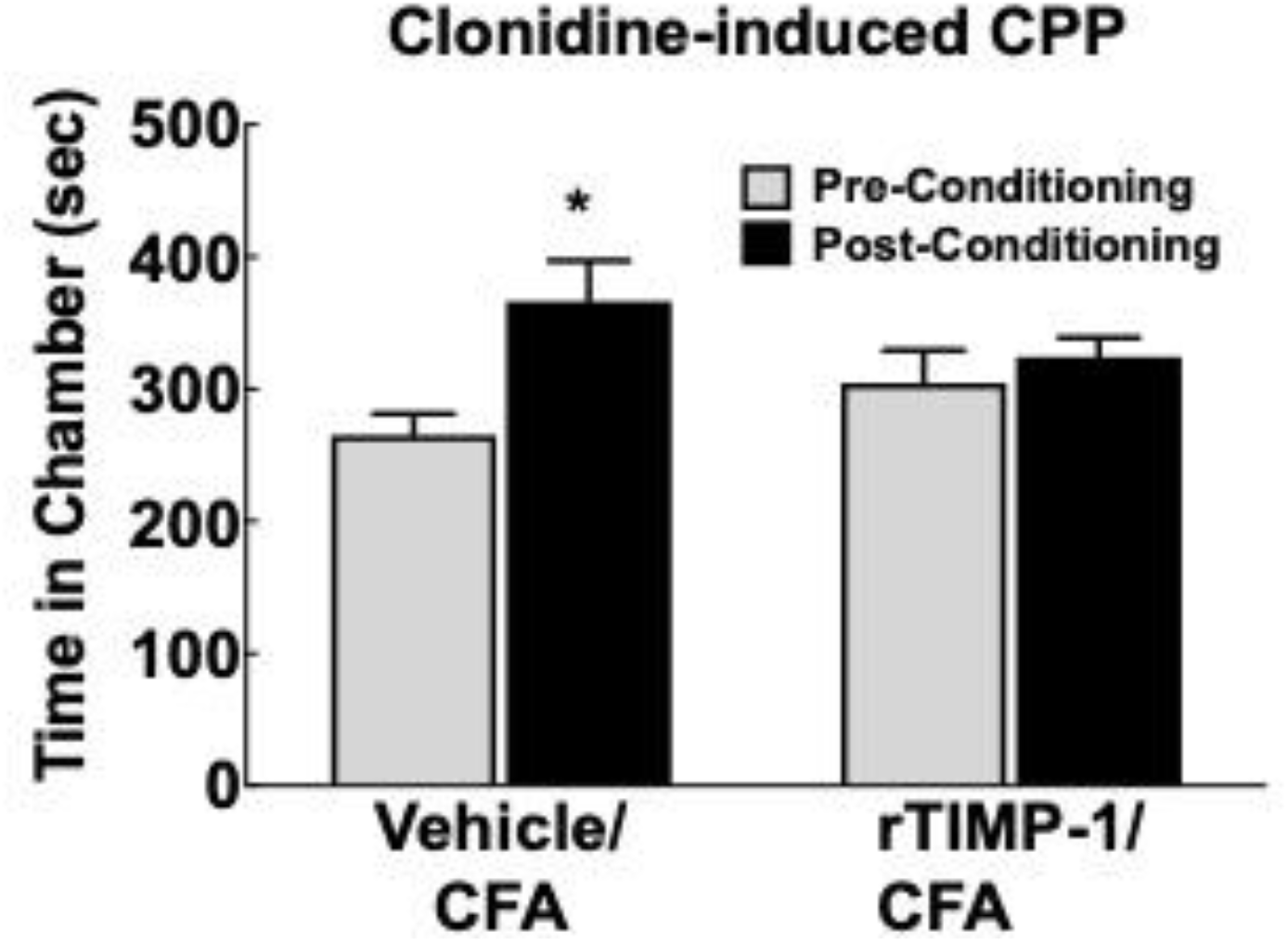
Replacement of TIMP-1 attenuates ongoing inflammatory pain in WT mice. Comparison of pre-conditioning and post-conditioning time spent in the clonidine paired chamber show a significant increase in the post-conditioning time in the Vehicle/CFA treated mice, but not the rTIMP-1/CFA treated mice. *p<0.01 vs pre-conditioning time. Sample size CFA/rTIMP-1 = 9; CFA/Veh = 8. Error bars = S.E.M.

## DISCUSSION

The balance between TIMPs and MMPs is important for maintaining tissue homeostasis and preventing pathological conditions. Following tissue damage and inflammation, TIMP-1 is expressed in a variety of cell types that can modulate neuronal function and wound healing, including, astrocytes, oligodendrocytes, Schwann cells, endothelial cells, mast cells, and keratinocytes (Fagerberg et al., 2010;Yokose et al., 2012;Claycomb et al., 2013). Because TIMP-1 is broadly expressed, TIMP-1 may also regulate neuroinflammation and neuropathic pain (Dhar et al., 2006;Kawasaki et al., 2008;Ji et al., 2009;Huang et al., 2011). Although the predominant view is that TIMP-1 exerts these functions by inhibiting MMPs, emerging evidence suggests that TIMP-1 may also facilitate these functions by binding cell-surface receptors and mediating their subsequent downstream signaling pathways (Dhar et al., 2006;Toricelli et al., 2013;Thevenard et al., 2014;Nicaise et al., 2019). Therefore, the present set of studies was designed to investigate the role of TIMP-1 in the development of inflammatory hypersensitivity and the mechanisms of its action.

To determine how inflammation affected the expression of TIMP-1 in tissues proximal and distal to the site of CFA-injection, we examined TIMP-1 expression in skin, DRG, and spinal cord over the course of 7 days. While we did not observe changes in TIMP-1 expression in the DRG or spinal cord, we did localize its expression to GFAP-positive cells, suggesting that TIMP-1 is expressed by satellite glial cells and astrocytes, respectively. When we examined skin, we found that CFA induced an 8.27 fold increase in TIMP-1 protein expression within 24 hr of inflammation, and that this temporal upregulation in TIMP-1 was observed in basal keratinocytes. Given that keratinocytes augment nociceptive signaling through the release of neuroactive molecules (Baumbauer et al., 2015;Moehring et al., 2018), the release of TIMP-1 from keratocytes may attenuate pronociceptive behavior caused by inflammation (Fagerberg et al., 2010;Yokose et al., 2012;Edqvist et al., 2015). To test this hypothesis, we examined the temporal expression pattern of TIMP-1 in relationship to the development of cutaneous hypersensitivity in WT mice. We found that the largest change in TIMP-1 expression, within 24 hr of injection, preceded the onset of cutaneous hypersensitivity, and that at 3 days when TIMP-1 expression peaked, behavioral sensitivity was the greatest in WT mice. This result suggests that peak TIMP-1 expression may in some way signal the onset of hypersensitivity. Alternatively, it is also possible that the relationship between TIMP-1 expression and the onset of hypersensitivity is determined by a relative change in expression between two consecutive time points. In the case of the current experiments the largest change in expression is observed between baseline and 24 hr following inflammation, where we observe an 8.27 fold change in expression. Conversely, we only detect a 1.43 fold increase in expression between Day 1 and Day 3, implying that it is not the absolute level of TIMP-1 expression, *per se*, that contributes to the delay in hypersensitivity, but rather the extent to which TIMP-1 expression changes relative to previous levels of expression over time. Consequently, it may be possible that as TIMP-1 expression increases during the first 24 hr of inflammation the emergence of pain-related behavior is attenuated. However, the overall change in TIMP-1 expression over the next 48 hr is no longer sufficient to prevent the emergence of hypersensitivity. Supporting this, we found that the replacement of recombinant TIMP-1 within 24 hr of CFA-injection, which causes a significant increase in expression from baseline expression in the skin, prevented the onset of hypersensitivity in mice lacking TIMP-1. Taken together, these data suggest that the immediate induction and release of TIMP-1 from basal keratinocytes attenuates inflammatory hypersensitivity.

If the release of TIMP-1 is important for delaying the onset of inflammatory hypersensitivity, it is also possible that hypersensitivity is exacerbated in the absence of TIMP-1. To test this, we compared mechanical hypersensitivity and thermal hyperalgesia at the site of inflammation in WT and T1KO mice. While we observed a robust reduction in mechanical thresholds in WT and T1KO mice, hypersensitivity persisted for a longer duration in T1KO mice relative to WT controls. Interestingly, when we examined thermal hyperalgesia, we observed a significant reduction in paw withdrawal thresholds in T1KO mice, while WT mice appeared to be unaffected. This result suggests that TIMP-1 may differentially regulate the processing of thermal and mechanical stimulation. In particular, in WT mice, the presence of TIMP-1 appears to delay the onset and persistence of mechanical sensitivity while having no effect on thermal reactivity during mild inflammation. More broadly, others have argued that mechanical sensitivity is a hallmark sign of pathological pain states (Treede et al., 1992;Treede et al., 2015), and our results imply that the dysregulation in TIMP-1 signaling may contribute to this process by influencing hypersensitivity to mechanical stimulation.

Given that thermal hyperalgesia and prolonged mechanical hypersensitivity were observed in T1KO, but not WT mice, following mild inflammation, our data suggest that the absence of TIMP-1 increases susceptibility to subtle perturbations that would otherwise be considered innocuous. Indeed, we found that injections of normal physiological saline produced mechanical hypersensitivity in T1KO mice, which was not observed in WT controls. This hypersensitivity could be due to a number of plausible factors, including hypertonicity (Steinbrocker et al., 1953;Giesler and Liebeskind, 1976;Hylden and Wilcox, 1983;Sutaeg Hwang and Wilcox, 1986), the disruption of cutaneous integrity from needle insertion, cutaneous distention following injection, or the induction of some inflammatory process. Whatever the cause of hypersensitivity is following saline injection, it is tempting to postulate that when TIMP-1 is not present, the physiological processes in the periphery become dysfunctional and sensory stimulation is amplified.

Pathological pain is also characterized by increased sensitivity in tissues adjacent to, and distal from, the site of inflammation, as is the case with “mirror image” pain (Shir and Seltzer, 1991;Maleki et al., 2000). When we assessed sensitivity in tissue adjacent to the site of inflammation and that are innervated by different sets of afferent terminal endings (e.g., hairy vs. glabrous skin), we found that T1KO mice exhibited mechanical hypersensitivity. Interestingly, we observed hypersensitivity in the uninflamed (contralateral) hindpaw, and the sensitivity occurred in a different dermatome from the inflamed dermatome. Administration of rmTIMP-1 at the site of inflammation alleviated hypersensitivity on both the inflamed and uninflamed paws. These data suggest that inflammation-induced TIMP-1 expression occurs in a coordinated fashion that influences the normal progression of inflammatory sensitivity in both inflamed and uninflamed tissues, which may attenuate afferent input and prevent the development of central sensitization.

TIMP-1 is well known as an inhibitor of MMPs, and we know that MMPs contribute to pain following various injury and inflammatory conditions (Kawasaki et al., 2008;Remacle et al., 2018). We hypothesized that disrupting the balance between TIMP and MMP expression and activity would exacerbate hypersensitivity in T1KO mice due to elevated MMP activity and pro-inflammatory cytokine expression. However, we did not detect any genotype-specific differences in the activity of MMP-9 or pro-inflammatory cytokines proximal to the site of CFA administration. Prior work indeed shows that the inhibition of MMPs reduces pain (Kawasaki et al., 2008;Ji et al., 2009), and our current results demonstrate that administration of the N-terminal domain of TIMP-1, the domain responsible for MMP inhibition, attenuated hypersensitivity. We also show that administration of TIMP-1(C), the domain responsible for engaging receptor-mediated cell signaling events (Jung et al., 2006;Toricelli et al., 2013;Takawale et al., 2017), also attenuates hypersensitivity following inflammation, suggesting that TIMP-1 may also delay the emergence of hypersensitivity through a novel receptor-mediated mechanism. Consequently, TIMP-1 may attenuate the development of pain through both pathways. This latter point may help to explain, at least in part, why small molecule inhibitors of MMP activity have limited efficacy (Cathcart and Cao, 2015). By understanding how both subdomains alleviate hypersensitivity, we may be able to effectively manage pain progression.

While our results show that TIMP-1 attenuates pain and hypersensitivity through both MMP inhibition and receptor-mediated signaling, the precise mechanisms by which TIMP-1 acts are not known. Our data suggest that the amount of TIMP-1 present at the site of inflammation may determine functional outcomes, which is consistent with previously published work (Grünwald et al., 2018). The function of TIMP-1 is also determined by the specific interactions TIMP-1 has with its binding partners, which includes both proteases and membrane-bound receptors. We have yet to identify which receptor is responsible for the antinociceptive properties of TIMP-1, but it is known that TIMP-1 binds and activates the CD63/β1 integrin receptor complex (Jung et al., 2006;Toricelli et al., 2013;Takawale et al., 2017;Nicaise et al., 2019). Interestingly, prior work has shown that interfering with the interaction between β1 integrin and versican attenuates inflammatory and neuropathic pain, as well as nociceptor activity(Dina et al., 2004;Ferrari and Levine, 2016). Consequently, keratinocyte-derived TIMP-1 may bind to CD63/β1 integrin expressed on cutaneous nerve endings ultimately, attenuating primary afferent function. While primary afferents are known to express β1 integrin, it is unclear whether they also express CD63. If neurons do not express CD63 there may be an indirect pathway involving other cell types, such as mast cells or keratinocytes, where TIMP-1 may bind to CD63 to alter the release pronociceptive molecules that influence neuronal function (Bertaux et al., 1991;Baumbauer et al., 2015;Moehring et al., 2018). Interestingly, disrupting the activity of β1 integrin prevents persistent pain in a model of hyperalgesic priming while leaving the acute phase of sensitivity intact(Dina et al., 2004). Therefore, stimulating TIMP-1 release, or the direct delivery of TIMP-1, may prevent the emergence of pathological pain, through β1 integrin activation, while leaving the capacity to detect normally painful stimuli unaffected.

Finally, demonstrating that TIMP-1 alleviates ongoing pain in WT mice we show that TIMP-1 is a potential clinical target for therapeutic intervention. It may be possible that TIMP-1 acts as a physiological “brake” on the nociceptive system to prevent overexcitation of primary afferents and the development of centralized pain states. Consequently, targeting TIMP-1 may have therapeutic benefit for both peripheral and central pain. For example, while we did not directly assess hypersensitivity at somatic regions beyond the contralateral hindpaw, our data may have implications for understanding the underlying mechanisms of widespread pain syndromes, such as fibromyalgia. Our data may also have implications for understanding metastatic processes in nonpainful forms of cancer. Indeed, TIMP-1 has been studied extensively in cancer (Gong et al., 2013;Toricelli et al., 2013;Jackson et al., 2017) and is therefore of significant interest for determining how metastases develop without producing pain. It is not yet clear whether painful and nonpainful cancers differentially express TIMP-1 or whether TIMP-1 receptor binding kinetics are altered in painful and non-painful cancers. Understanding these dynamics may have clinical implications for developing early cancer detection strategies, especially if TIMP-1 is considered a regulator of pain state and not just one involved in tissue remodeling. Finally, if TIMP-1 can be utilized as a target for attenuating pathological pain, TIMP-1 may serve as an alternative to opioid-based medicines. This possibility is intriguing given our data showing that peripheral administration of TIMP-1 has antinociceptive properties and prevents the spread of sensitivity to uninflamed somatic regions. However, these conclusions should be taken with some caution as research has shown that excess TIMP-1 expression, specifically through genetic overexpression, may lead to unintended adverse events (Grünwald et al., 2018). Therefore, future research should focus on illuminating the mechanisms by which TIMP-1 attenuates pain and hypersensitivity and how targeting this system may be therapeutically beneficial.

## ACKNOWLEDGEMENTS

Research was supported by the National MS Society grant RG-1802-30211(SJC).

We would like to thank Ms. Nicole Glidden and Ms. Jessica Yasko for their technical assistance.

